# Frequent and Transient Acquisition of Pluripotency During Somatic Cell Trans-Differentiation with iPSC Reprogramming Factors

**DOI:** 10.1101/008284

**Authors:** Itay Maza, Inbal Caspi, Sergey Viukov, Yoach Rais, Asaf Zviran, Shay Geula, Vladislav Krupalnik, Mirie Zerbib, Rada Massarwa, Noa Novershtern, Jacob H. Hanna

**Author notes:** Correspondence should be addressed to Jacob H. Hanna.

## Abstract

Recent reports have proposed a new paradigm for obtaining mature somatic cell types from fibroblasts without going through a pluripotent state, by briefly expressing canonical iPSC reprogramming factors *Oct4*, *Sox2*, *Klf4*, *c-Myc* (abbreviated as OSKM) in cells expanded in lineage differentiation promoting conditions. Here we apply genetic lineage tracing for endogenous Nanog locus and X chromosome reactivation during OSKM induced trans-differentiation, as these molecular events mark final stages for acquisition of induced pluripotency. Remarkably, the majority of reprogrammed cardiomyocytes or neural stem cells derived from mouse fibroblasts via OSKM mediated trans-differentiation (∼>90%), are attained after transient acquisition of pluripotency, and followed by rapid differentiation. Our findings underscore a molecular and functional coupling between inducing pluripotency and obtaining “trans-differentiated” somatic cells via OSKM induction, and have implications on defining molecular trajectories assumed during different cell reprogramming methods.

## Main Text

The identity of somatic cells can be converted to variant extents by ectopic expression of exogenous factors^1,2^. The latter include the ability to directly reprogram somatic cells to pluripotency via ectopic expression of exogenous transcription factors, canonically OSKM, in Leukemia Inhibitory factor (LIF) containing mouse ES growth conditions^2^. Isolated iPSCs (OSKM-iPSC) can be subsequently directed towards a variety of lineages by distinct differentiation protocols. Since their initial discovery, a variety of factors can substitute or complement OSKM components to yield iPSCs at greater efficiencies and kinetics^3,4^. Importantly, cytokines and small molecules used can directly influence epigenetic reprogramming by modulating endogenous signaling and chromatin remodeling pathways (e.g. LIF/Stat3 signaling pathway activation, Vitamin C induced epigenetic changes, BMP4 promotion of naïve pluripotency, TCF3 target gene derepression following GSK3 inhibition etc.)^5,6^.

A different strategy for lineage conversion, termed somatic cell trans-differentiation, entails ectopic expression of somatic lineage master regulators that induce mature somatic cells cell transformation into a different cell type without going through a pluripotent configuration (e.g. MyoD converts fibroblasts into muscle cells, C/EBPα converts Pro-B cells into macrophage-like cells, Mitf converts fibroblasts into melanocyte-like cells)^7–9^. Recently however, a new approach combining pluripotency induction and somatic trans-differentiation, called OSKM mediated trans-differentiation (OSKM-TD), has been introduced as a mean to directly interconvert somatic cells without acquiring pluripotency^10–12^. In this approach, instead of applying defined somatic lineage transcription factors, the four Yamanaka pluripotency reprogramming factors are briefly induced to periods as short as 3-10 days to presumably only induce a partially reprogrammed plastic state^10,11^. By providing lineage-specifying media that does not contain conventional pluripotency promoting cytokines like LIF, reprogramming of such intermediate supposedly “plastic” state is shifted towards a desired somatic cell fate, without attaining a state of pluripotency. Notably, by using this approach to achieve somatic cell trans-differentiation, typically higher quality and more fully programmed cells are produced, as compared to those achieved via ectopic expression of somatic lineage promoting factors^11^.

Several experimental pillars were provided to exclude acquisition of pluripotency in the OSKM-TD approach^10–12^. First, brief OSKM induction of only up to maximum of 10 days was applied, which was deemed as insufficient to yield iPSCs^10,11^. Further, these studies reported failure to detect Nanog-GFP expression or transcription in bulk cultures assayed at different time points during OSKM-TD^10,11^. Absence of LIF and utilization of JAK1 small molecule inhibitors to block LIF/JAK1/Stat3 signaling were also used to exclude acquisition of pluripotency. Despite of the above, lineage tracing tools to unequivocally exclude acquisition of pluripotency during this manipulation in the rare cells that successfully trans-differentiate among the donor cell population (reported efficiencies <1%), were not used^10–12^. Thus it remains unclear whether somatic cells produced by this technique are in fact trans-differentiated or rather preferentially transiently go through a state of induced pluripotency and then differentiate to a somatic lineage according to the differentiation media conditions used in OSKM-TD protocols.

Our interest to carefully examine the OSKM trans-differentiation phenomena was evoked via multiple results that strengthened the possibility the iPSCs may still be formed under suboptimal reprogramming conditions, and that brief OSKM induction can overcome differentiation signals and leads to rapid iPSC formation. First, we were able to obtain Nanog-GFP+ iPSCs at low efficiency from different Doxycycline (Dox) inducible OSKM transgenic systems with as little as 3 days of Dox induction (Fig. 1a). These results are consistent with previous reports that obtained iPSC after brief OSKM induction and existence of privileged donor somatic cells in fibroblasts that rapidly commit and convert into iPSC^13–15^. Further, when *Oct4-GFP* reporter fibroblast cells were subjected to OSKM transduction in Cardiogenic or neural stem cells (NSCs) growth conditions, instead of pluripotency conditions, we were still able to obtain GFP expression ES-like colonies and observed “hybrid” colonies with Oct4-GFP+ cells in the center of the colony while their edges showed clear neuronal differentiation signs and lack of Oct4-GFP **(Supplementary Fig. 1)**. The results above emphasized the need to exclude the possibility that iPSCs may still be formed under suboptimal reprogramming conditions and may be a considerable source for mature trans-differentiated cells obtained via OSKM-TD approaches^10,11^.

**Figure 1:**
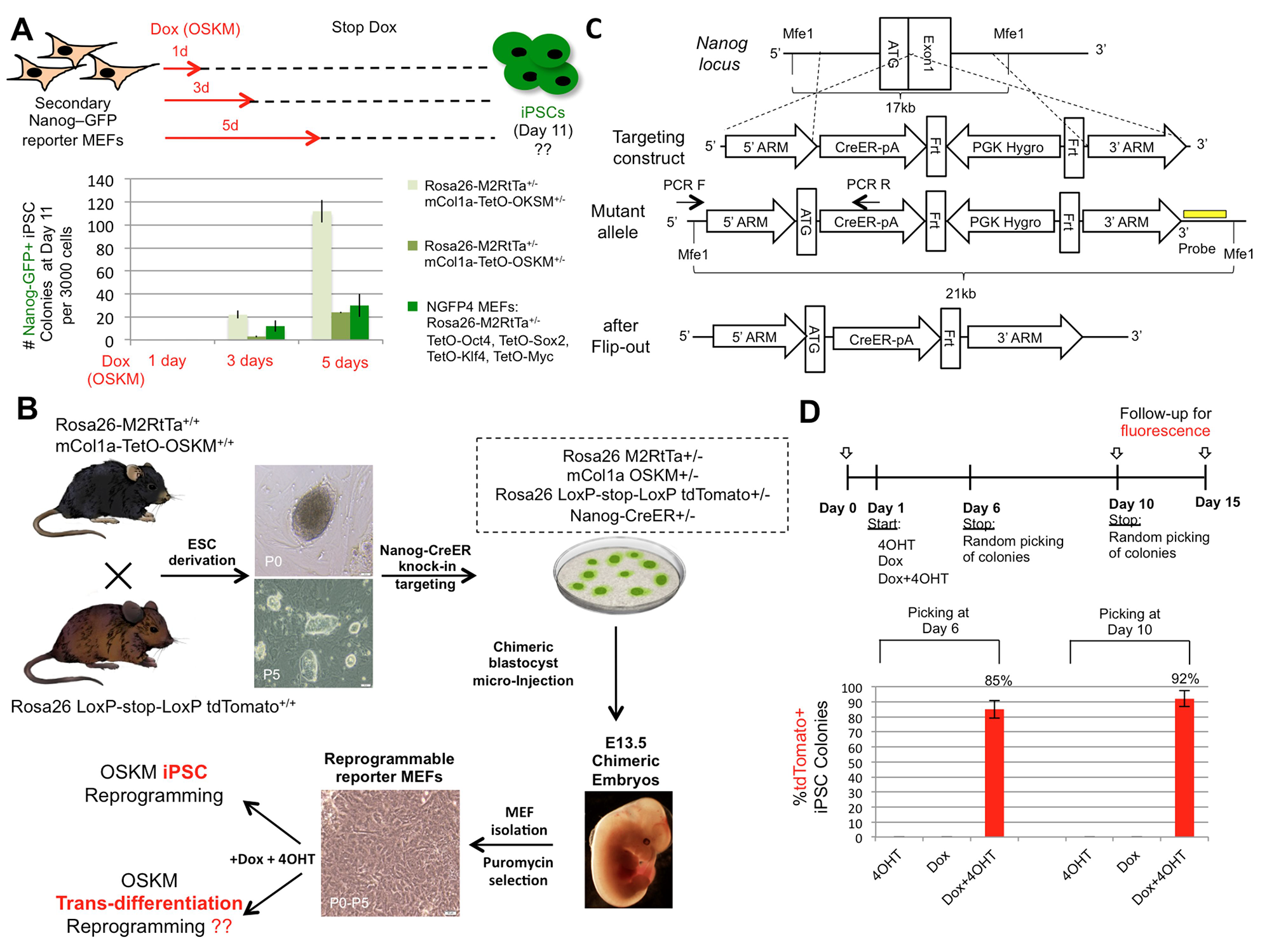
Lineage tracing for endogenous Nanog reactivation during reprogramming. **A.** Secondary MEF cells from three indicated different secondary reprogramming systems and carrying Nanog-GFP knock-in reporter for pluripotency, were subjected to Dox induced reprogramming. Dox was applied for the indicated time points, and then withdrawn. iPSC formation was evaluated at day 11 without passaging. Error bars indicate SD of biological duplicates (1 out of 2 representative experiments is shown). **B.** Scheme illustrating generation of Quadruple knock in allele reporter reprogrammable MEFs utilized for either OSKM-iPSCs or OSKM trans-differentiation (OSKM-TD) reprogramming. **C.** Simplified schematic illustration of the Nanog-CreER knock in targeting construct, directed to the endogenous Nanog locus. **D.** Reprogrammable Nanog-CreER MEFs were subjected to iPSC reprogramming protocol in the presence of Dox, 4OHT or both, which were withdraw at day 6 or 10 as indicated. 48 Colonies were picked and validated as iPSCs (Nanog/ SSEA1 and AP staining) from each condition, and then scored for the presence of tdTomato+ signal. Bar plot indicating % of tdTomato+ iPSC colonies obtained, indicating the sensitivity of the system. Single colonies were randomly subcloned and examined for tdTomato fluorescence at two time points day 6 and day 10, revealing average sensitivity (fidelity) of 85-92%, respectively. In the absence of 4OHT, none of the iPSCs had tdTomato+ signal, indicating 100% specificity of the system. Error bars indicate SD of biological duplicates (1 out of 3 representative experiments is shown).

We next moved to engineer an exacting system to track transient acquisition of pluripotency during in vitro reprogramming. We aimed to design a genetic tracing system for endogenous *Nanog* reactivation, as it is an important factor for establishing pluripotency *in vivo* and *in vitro*, as its reactivation occurs at the late and final stages of iPSC formation in mouse and human cells^10,11,16,17^. Consistently, the previous OSKM-TD studies heavily focused on excluding *Nanog* reactivation and transcriptional detection during OSKM mediated trans-differentiation in the bulk reprogramming cultures^10,11^ as conclusive evidence for excluding acquisition of pluripotency. We designed a reprogrammable system in which transient activation of *Nanog* in the process of reprogramming can be easily monitored by permanent tdTomato fluorescence activation, indicating that pluripotency was achieved, even if only temporarily. We first cross-bred *Rosa26 RtTa/m.Col1a OSKM^+/+^*reprogrammable mouse strain^18^ with *Rosa26 LoxP-stop-LoxP tdTomato^+/+^ reporter mice, and* derived and validated ESC lines from day E3.5 blastocysts that are heterozygous for the triple knock-in alleles: *Rosa26 M2RtTa^+/-^, Rosa26 LoxP-Stop-LoxP tdTomato^+/-^*and *m.Col1a-TetO-OSKM ^+/-^* (Fig. 1B). Subsequently, we generated a BAC recombineered Nanog-CreER targeting construct, which introduces a knock-in Tamoxifen inducible CreER cassette under the control of endogenous Nanog promoter (Fig. 1C). The quadruple knock-in ESC line was subcloned after PCR and Southern Blot validation for correct targeting by Nanog-CreER knock in allele **(Supplementary Fig 2A.)**. The latter ESCs were microinjected in host blastocyst to generate chimeric animals, from which transgenic fibroblasts were extracted for experimental analysis (Fig. 1B).

We validated the sensitivity and specificity of our system. Mouse ESCs turned on tdTomato rapidly after 48 hours of 4OHT induction (100% of ES colonies), while no fluorescence was detected in ESCs passaged over extended time in the absence of 4OHT **(Supplementary Fig 2B-C)**. MEF cells expanded with or without Dox in the absence of 4OHT, did not show any reactivation of tdTomato, further indicating 100% specificity of the system and excluding leakiness during expansion or reprogramming (Fig. 1D). The fidelity (sensitivity) of the reporter system was evaluated by reprogramming the transgenic MEFs to pluripotency by adding Dox and 4OHT to the medium and withdrawing both molecules at two time points: 6, and 10 days. Adding 4OHT alone did not produce any colonies, while Dox alone did not produce any tdTomato+ colonies (Fig. 1D). Formed ES-like colonies for each of the conditions, at the abovementioned time points (48 colonies for each group), were randomly subcloned and grown for another 5-6 days at which point the presence of tdTomato+ clones was assessed as a percentage of the total number of developed validated iPSC clones (AP+/OCT4+/SSEA1+ independent of Dox). We found the fidelity of our system to be ∼85% and ∼92% for iPSCs obtained after 6 and 10 days of Dox induction, respectively (Fig. 1D). Lack of 100% sensitivity might be explained by a number of non-mutually exclusive possibilities: limitations of the Nanog knock-in allele activation, mono-allelic expression of Nanog^19^, or reflect true biological outcome particularly that Dox/4OHT pulses are given for a limited period of time during reprogramming (only 6 or 10 days). Nevertheless, the above results validate the ability of the system to specifically document authentic acquisition of pluripotency during iPSCs reprogramming and with very high sensitivity.

We subjected *Nanog* tracing system and reporter engineered MEF cells to previously reported OSKM trans-differentiation experiments converting fibroblasts into cardiomyocytes or neural stem cells (NSCs). Importantly, protocols of experiments and media used were meticulously followed as published in the relevant papers^10,11^. Direct reprogramming of *Nanog-CreER* reporter MEFs to NSCs was done in knockout DMEM medium with 10% KSR/5% FBS and with Dox and 4OHT for the first 3-6 days (Fig. 1A). Thereafter, Dox and 4OHT were withdrawn from the medium, and cells were grown in neural reprogramming medium (DMEM-F12 and Neurobasal medium mixed by 1:1 and supplemented with FGF2 and EGF) until NSC colonies were formed by day 13 (Fig. 1B). From day 4 onwards JAK1 inhibitor (J1I) was added to the culture medium. At day 13, formed colonies were subcloned and grown in NSC medium on Poly-D-Lysine and Laminin coated plates. All experiments of OSKM mediated trans-differentiation of MEFs into cardiomyocytes or NPCs were carried out in LIF-free mediums, and without the presence of feeder cells. Indeed Sox1+ NSC colonies were apparent in this protocol, and could be subcloned expanded as stable line, and were expectedly responsive for induced differentiation into differentiated Tuj1+ neurons **(Supplementary Fig. 3A-B)**. RNA-seq analysis confirmed their transcriptional similarity to primary embryo derived NSCs, and their distinct form mouse fibroblasts or ESCs **(Supplementary Fig. 3C)**. Remarkably, ∼88% of Sox1+ NPC colonies were tdTomato+ indicating acquisition of pluripotency during OSKM mediated somatic cell conversion

Conversion of MEFs into cardiomyocytes was done in reprogramming media containing 15% fetal bovine serum (FBS) and 5% knock-out serum replacement (KSR) with added JAK1 inhibitor (JI1) for 6 days, followed by switch to 1% FBS and 14% KSR and JAK1 inhibitor for 3 days (Fig. 2D). Dox and 4OHT were added to the medium at day 1 and continuously kept in the medium until day 6-9. From day 9 onwards, cells were cultured in chemically defined RPMI/N2B27 medium, with the cardio-inductive growth factor BMP4 for the first 5 days (Fig. 2D). Colonies were allowed to develop for up to 21 days or until beating cardiomyocytes were noticed in the plated cells. Indeed by applying this protocol we were able to achieve cardiomyocytes that expressed early and late cardiomyocytes markers (*Myosin, cardiac Troponin)* many of which showed beating rhythmicity (Fig. 2f ***and Supplementary Fig. 3D)***. Remarkably however, ∼93% of Myosin positive cardiomyocytes colonies were tdTomato+ indicating that they reactivated pluripotency during their conversion (Fig. 2E-F, ***Supplementary Fig. 3E***, **Supplementary Videos 1-6**).

**Figure 2:**
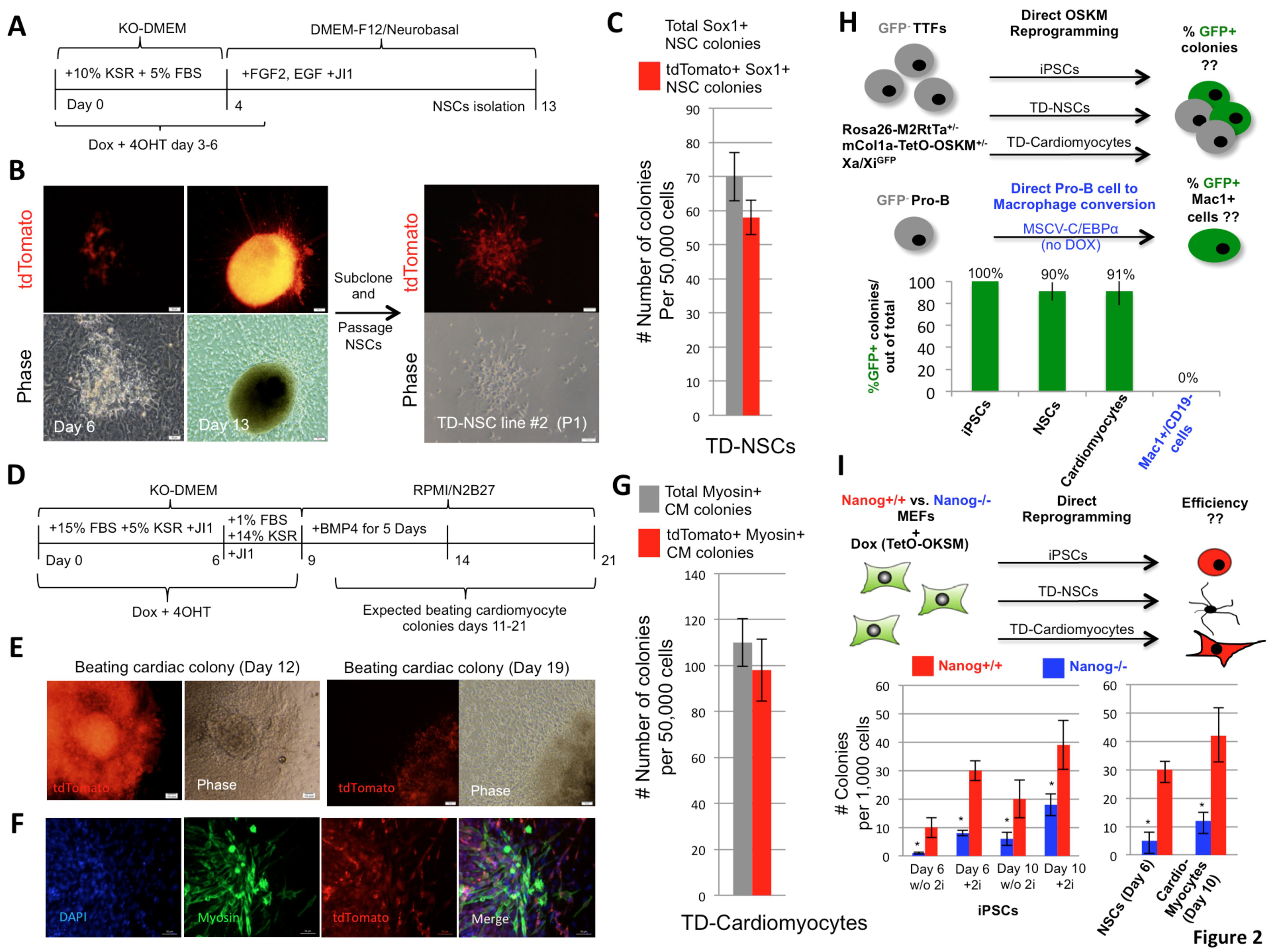
Frequent and transient acquisition of pluripotency during OSKM-TD. **A.** Schematic illustration of NSCs trans-differentiation with indicated time points for medium switch, small molecules and growth factors additions. **B.** Representative images of NSCs tdTomato+ colonies generated by the above mentioned protocol (**A**), at the start and end points of neural reprogramming. Passage of subcloned NSCs colony grown in NSCs medium, showing typical NSCs morphology and tdTomato+ fluorescence. **C.** Bar plot of immunostained Sox1+ TD-NSC colonies, showing the number of total and of tdTomato+ colonies obtained. Error bars indicate SD of biological triplicate wells (1 out of 2 representative experiments is shown). **D.** Schematic illustration of cardiomyocytes trans-differentiation with indicated time points for medium switch, small molecules and growth factors additions. **E**. Representative images of cardiomyocytes colonies reprogrammed by the above-mentioned protocol (**D**) with tdTomato+ fluorescence, and repeated rhythmic beating motion after 12 or 19 days of reprogramming. **F**. Representative immunostaining for Myosin in trans-differentiated cardiomyocytes colonies with co-localization of tdTomato fluorescence. **G**. Bar plot quantifying validated immunostained Myosin+ trans-differentiated cardiomyocytes colonies, showing total number and of tdTomato+ colonies obtained. Error bars indicate SD of biological triplicate wells (1 out of 2 representative experiments is shown). **H.** Tail tip fibroblasts (TTFs) were established from the indicated adult transgenic reprogrammable female mice. GFP-populations were sorted and purified and subjected to OSKM iPSC or TD protocols. Pro-B cells were purified form the same mice, and subjected to C/EBPa mediated conversion to macrophage like cells (without OSKM induction). Percentage of GFP+ cells or GFP+ cell containing colonies detected at day 13 (for OSKM reprogramming) and day 8 (for Pro-B to macrophage conversion) is indicated. Error bars indicate SD of biological duplicates (1 out of 2 representative experiments is shown). **I.** Reprogramming efficiencies of Nanog^-/-^ and Nanog^+/+^ MEFs, to either iPSCs, NSCs or cardiomyocytes, evaluated by alkaline phosphatase (AP), Sox1 or Myosin colony staining, respectively. Reprogramming was conducted following transduction with FUW-M2RtTa and FUW-TetO-OKSM lentiviral vectors. Error bars indicate SD of biological triplicates (1 out of 2 representative experiments is shown). *Indicates student t-test P value <0.05 (in comparison to matched Nanog^+/+^ sample). Bar plot of colony number indicates reduction in reprogramming efficiency of Nanog^-/-^ MEFs, as dependent on duration and reprogramming conditions.

We next aimed to apply a different lineage tracing system for evaluating acquisition of pluripotency during OSKM-TD, by determining whether the inactive X chromosome in donor female somatic cell lines undergoes reactivation. It has been well established that X chromosome reactivation is a late and specific hallmark feature for reactivation of pluripotency and is followed by random X inactivation in clonal iPSC derived differentiated progenies^20^. To track X chromosome reactivation and inactivation dynamics, we crossed female *Rosa26 M2RtTa^+/+^* and *m.Col1a-TetO-OSKM ^+/+^ mice*^18^ with male mice carrying an X-linked GFP reporter transgene^21^, and purified GFP-tail tip fibroblasts form female mice (i.e., all GFP-reprogrammable cells carry the GFP transgene on the inactive X chromosome) (Fig. 2H). Remarkably, GFP fluorescence was identified in >90% of OSKM-TD NSC and cardiac colonies (Fig. 2H), and in 100% of iPSCs obtained. Importantly, reprogramming of female derived GFP-Pro-B lymphocytes into macrophage like cells by C/EBPα overexpression^22^, did not yield any GFP+/Mac+ colonies (Fig. 2H), excluding promiscuous and non-specific X chromosome reactivation in any transdifferntiation protocol applied. The latter results reinforce Nanog reactivation lineage tracing results (Fig.2 A-G), and reiterate the conclusion that the vast majority of OSKM-TD derived cells pass through a pluripotent state before transitioning towards somatic differentiation.

In the next set of experiments, we set out to explore a functional role for endogenous *Nanog* reactivation in OSKM mediated trans-differentiation, as the latter event has been shown not only to mark final stages of pluripotency acquisition, but to also synergistically boost iPSC reprogramming by OSKM^23^. We utilized previously described *Nanog^-/-^* knock out and *Nanog^+/+^* ESCs^23^ and as recently published, Nanog^-/-^ ESCs can be expanded in 2i/LIF/15% KSR enriched conditions while maintaining their pluripotency^24,25^ (**Supplementary Fig. 4A**). Reprogramming of *Nanog^-/-^*MEFs and secondary *Nanog* wild type MEFs to pluripotency was done following TetO-OSKM transduction and plating in two triplicates of 1＾10^3^ cells plated on day 0 for each time point and medium condition, in a 6 well plate, followed by adding Dox at day 1 to the either with or without 2i supplement. Formed colonies at day 6 and 10 were evaluated and counted by alkaline phosphatase staining. Reprogramming efficiency to pluripotency of *Nanog^-/-^*MEFs as compared to *Nanog* wild type showed a reduction of 90% and 55% in efficiency at days 6 and 10, respectively (Fig. 2I **and Supplementary Fig. 4B-D)**. Next, *Nanog^-/-^* and *Nanog^+/+^* MEFs were used in OSKM mediated trans-differentiation of NSCs and cardiomyocytes. The efficiency of trans-differentiation was evaluated as the number of colonies stained positive for NSCs or cardiomyocytes markers respectively, relative to the total number of colonies emerged after OSKM induction. Experiments were carried out in two triplicates of 1＾10^3^ cells plated at day 0, in a 6 well plate, revealing a reduction of efficiency by 80-85% in the *Nanog^-/-^* MEFs as compared to *Nanog^+/+^*wild type cells (Fig. 2I **and Supplementary Fig. 4D)**. Collectively, these results show that endogenous Nanog reactivation is functionally coupled to OSKM-Transdifferntiation efficiency, and at an equivalent robustness to that observed during OSKM-iPSC formation (Fig. 2I **and Supplementary Fig. 4D)**.

Finally, OSKM mediated trans-differentiation regimens make use of mediums devoid of LIF or with added JAK-Inhibitor, which block Stat3 pathway signaling, assuming these conditions completely prevent the formation of pluripotent iPSCs^10,11^. Given that induced reprogramming OSKM transgenes has been previously found sufficient to support expansion of pluripotent cells form non-permissive strains like NOD in LIF only conditions^26^, and that they also boost EpiSC reprogramming^24^, we wondered if they can completely supplement for exogenous LIF/Stat3 signaling. To tackle the latter issue, reprogramming of *Nanog CreER* reporter MEFs to pluripotency was done with added JAK1-Inhibitor and LIF-free N2B27 medium (**Supplementary Fig. 5A**). Transgene dependent iPSCs reprogrammed in these conditions were derived, expanded and shown to possess bona fide pluripotent capacity as evident by, positive staining for alkaline phosphatase, SSEA1, Nanog and mature teratoma formation **(Supplementary Fig. 5A-D)**. These results corroborate previous finding that Stat3 can be dispensable for re-establishing pluripotency^6^, and that exclusion of LIF/Stat3 is not an unequivocal proof for lack of acquisition of pluripotency when providing exogenous pluripotency factor transgenes. It is also important to note that a variety of ingredients claimed to increase trans-differentiation efficiency, are well known boosters for iPSC induction and naïve pluripotency maintenance in mice, including BMP4 cytokine^27^ and Knockout serum replacement (KSR) that contains L-ascorbic acid (Vitamin C)^5,28^.

Conversion of cell fate *in vitro* or *in vivo* carries great potential for the future of regenerative medicine. While there are often different routes of reprogramming to achieve a certain defined cell type, each might carry a variety of technical and functional advantages and disadvantages that are of importance to be explored^29^. Simultaneously, it is critical to study the molecular paths and trajectories assumed by these reprogrammed cells^30,31^. As such, we revisited recently described methods to convert somatic cell types by canonical iPSC reprogramming factor expression^11,32^, and concluded that while the technique of OSKM mediated trans-differentiation may be a relevant and safe way to fully reprogram differentiated cells, most of the latter cells are obtained following temporarily reaching a pluripotent state and rapidly differentiating according to the media used. While our study cannot absolutely exclude that some trans-differentiated cells are assumed without going through pluripotency, it does indicate that vast majority of converted cells are preferentially derived via acquisition of pluripotency when OSKM are used. Finally, considering the increase in identification of alternative factors that can induce and maintain pluripotency^3,4,33^, our findings underscore the importance of utilizing unbiased molecular lineage tracing during cell reprogramming *in vitro* to track trajectories assumed by cells during different direct conversion approaches.

## Author Contributions

I.M, N.N. and J.H.H conceived the idea for this project. I.M. designed and conducted experiments and wrote the manuscript with contributions from other authors. M.Z. conducted microinjections. R.M. assisted in microscopy imaging. S.G., I.C., V.K. assisted in tissue culture and reprogramming experiments. S.V., Y.R. and I.M. constructed and targeted Nanog-Cre ER knock-in construct. A.Z. and N.N. conducted RNA-seq analysis.

## Acknowledgements

J.H. is supported by a generous gift from Ilana and Pascal Mantoux; a Robertson Innovator grant by the New York Stem Cell Foundation (NYSCF), a research grant from the ERC starting investigator program (StG-2011-281906), the BIRAX initiative, the Leona M. and Harry B. Helmsley Charitable Trust, The Sir Charles Clore Research Prize, the Israel Science Foundation (Bikura, ICORE and Regular research program), the ICRF Foundation, the Benoziyo Endowment fund, Fritz Thyssen Stiftung, the Alon Foundation scholar award, a grant from Erica A. and Robert Drake. We thank the Weizmann Institute management for providing critical financial and infrastructural support.

## Supplementary Figure Legends

**Supplementary Figure 1: Generation of Oct4-GFP+ colonies under cardiac and neural growth conditions. A.** Schematic representation of experimental reprogramming conducted.. TetO-OSKM transgenic secondary MEFs carrying Oct4-GFP knock in reporter were subjected to Dox reprogramming for 21 days in cardiomyocyte of NSC growth conditions without passaging. **B.** Representative images of Oct4-GFP+ colonies obtained from MEFs reprogrammed for 21 days on Dox (OSKM) induction in cardiomyocytes growth conditions **C.** Representative “Hybrid” colonies following reprogramming of transgene Oct4-GFP MEFs in NSC growth conditions, showing differentiated neurons expressing Tuj1 on the periphery of the colony, with transgene Oct4-GFP positive cells at the center of the colony. Note expected lack of co-localization between Tuj1+ and Oct4-GFP+ cells.

**Supplementary Figure 2: Nanog-CreER reporter system for endogenous Nanog locus reactivation. A.** Detailed schematic illustration of the mouse Nanog locus and Nanog-CreER knock in targeting construct, directed to the endogenous Nanog locus. Southern blot verification of Nanog Cre-ER knock-in to the triple allele ESCs is also shown. Both clones 1 and 2 showed correct targeting, and clone 1 was further chose for Flip-out and subsequent experimental lineage tracing analysis. **B.** Representative images of Nanog Cre-ER ESCs showing tdTomato activation when exposed to 4OHT for 48 hours (100% of colonies) and absence of fluorescence when grown with ETOH alone after 7 days. **C.** Representative images of reporter MEFs when exposed to 4OHT, Dox, or both showing no detection of colonies in MEFs exposed to 4OHT alone, no detection of tdTomato+ colonies in reporter MEFs grown with Dox alone, and tdTomato+ colonies in iPSC colonies obtained with Dox+4OHT but no detection of tdTomato in MEFs that did not undergo reprogramming under Dox+4OHT. The results show absence of leakiness of the system and specificity for Nanog-tdTomato reactivation only in ES-like colonies.

**Supplementary Figure 3: Validation and characterization of OSKM-TD cells. A.** Immunostaining of NSC markers: Nestin, Sox1 and Sox2 in primary brain derived NSCs (control Brain-NSC line #1 shown as a representative line at the indicated passage number), and trans-differentiated NSCs (TD-NSC line #1). Merged images of trans-differentiated NSCs indicating co-localization of tdTomato reporter with NSC markers. **B.** Differentiation of OSKM-TD NSCs into Tuj1 mature neurons. Tuj1 signal co-localizes with tdTomato as the latter label is expected to be permanent once turned on **C.** Hierarchical clustering heat map comparison of genetically matched mouse samples including primary brain derived NSCs, MEF, ESC and two subclones of trans-differentiated NSCs. The color bar indicates normalized gene expression, where red indicates genes that are over-represented in the specific sample. Values are in standard deviations. D. Immunostaining for cardiac specific troponin (cTnT) marker on beating OSKM-TD cardiomyocytes colony (validated as beating before fixation). Merged images of trans-differentiated cardiac cells indicating co-localization of tdTomato reporter with cTnT marker. **E.** Bar plot indicating percentage of tdTomato+ colonies out of the total number of OSKM trans-differentiated stained and validated NSCs and cardiomyocytes colonies. Colonies were indicated NSCs if immunostained positive for Sox1, and as cardiomyocytes if immunostained positive for Myosin/cTnT. Error bars indicate SD of biological triplicate wells (1 out of 2 representative experiments is shown).

**Supplementary Figure 4: Endogenous Nanog reactivation functionally support OSKM-TD reprogramming efficiency outcome. A.** Validation of Nanog-/-ESCs by western blot and immunostaining for Nanog, Oct4 and SSEA1. Nanog-/-ESCs express pluripotency markers Oct4, SSEA1, but do not express Nanog. **B.** Pluripotency validation of Nanog-/-iPSCs reprogrammed from Nanog-/- MEFs following OSKM induction, by immunostaining for pluripotency markers SSEA1 and Oct4 and alkaline phosphatase stain. **C.** Evaluation of pluripotency potential of transgene (Dox) independent Nanog-/- iPSCs, evident by teratoma formation assay. Images of Hematoxylin & Eosin stain of teratoma segments with representation of all three germ layers: endoderm (glandular epithelium), mesoderm (cartilage) and ectoderm (squamous epithelium). **D.** Reduction of reprogramming efficiencies of Nanog-/- MEFs to iPSCs, NSCs and cardiomyocytes as compared to control Nanog+/+ MEFs. Bar plot indicates ratio reduction of reprogramming efficiency, evaluated by number of iPSC colonies per 1,000 plated cells. Noticeably, reprogramming efficiency was similarly reduced in Nanog-/- MEFs when reprogrammed to iPSC without 2i at day 6 and 10, compared to Nanog-/- MEFs reprogrammed to NSCs (quantified after 6 days Dox activation) and cardiomyocytes (quantified after 10 days Dox activation).

**Supplementary Figure 5: OSKM exogenous transgenes maintain iPSC pluripotency independent of exogenous LIF/Stat3 signaling. A.** Schematic representation of reporter MEFs reprogramming to iPSC with JAK1 inhibitor (JAK1I) and exogenous LIF free conditions. **B.** Representative images of reprogrammed reporter MEFs at day 5 reprogramming and established clones with or without Dox maintained in the growth medium, with activation of tdTomato Nanog-CreER reporter. At P1, the colonies differentiated after Dox withdrawal (upper panel), ES-like colonies can be maintained on Dox (lower panel). **C.** Alkaline phosphatase stain and Immunostaining for pluripotency markers SSEA1 and Nanog in iPSC reprogrammed in JAK1 inhibitor and LIF free conditions from reporter MEFs, showing co-localization with tdTomato. **D.** Evaluation of pluripotency potential of iPSCs reprogrammed with JAK1 inhibitor and in LIF free conditions from reporter MEFs, as evident by teratoma formation assay. Images of Hematoxylin & Eosin stain of teratoma segments with representation of all three germ layers: endoderm (glandular epithelium), mesoderm (cartilage) and ectoderm (squamous epithelium). The results validate induction and maintenance of bona fide pluripotency in mouse iPSCs by exogenous OSKM transgene expression and without LIF/Stat3 signaling.

### Supplementary Video legends

**Supplementary Video 1**: Example #1: Phase-contrast live imaging of a forcefully contracting OSKM-TD cardiomyocyte colony.

**Supplementary Video 2**: Example #1: tdTomato fluorescent live imaging of the beating cardiomyocyte colony shown in Supplementary Video 1.

**Supplementary Video 3**: Example #2: Phase-contrast live imaging of a forcefully contracting OSKM-TD cardiomyocyte colony.

**Supplementary Video 4**: Example #2: tdTomato fluorescent live imaging of the beating cardiomyocyte colony shown in Supplementary Video 3.

**Supplementary Video 5**: Example #3: Phase-contrast live imaging of a forcefully contracting OSKM-TD cardiomyocyte colony.

**Supplementary Video 6**: Example #3: tdTomato fluorescent live imaging of the beating cardiomyocyte colony shown in Supplementary Video 5.

## Experimental procedures

### Generation of Reprogrammable Nanog-CreER Reporter Mice

Homozygote *Rosa26 LoxP-stop-LoxP tdTomato^+/+^* mice were a kind gift by Prof. Eliezer Zelzer (Weizmann Institute of Science). These mice were crossbred with the reprogrammable double homozygote *Rosa26-M2RtTa* and *m.Col1a OSKM^+/+^* mice^18^. At day E3.5 blastocysts were extracted and individually seeded in a flat bottom 96 well with plated gamma-irradiated feeder cells, in mESC growth medium: DMEM (*gibco 41965-039*) supplemented with human LIF (20ng/ml prepared in house), 1% L-Glutamine, 1% Non-essential amino acids, 1% penicillin/streptomycin, 0.1 mM β-mercaptoethanol and 15% FBS (Biological Industries). Developed mESCs colonies were expanded as single clonal lines. In order to make a targeting construct for inserting CreERT2 gene in frame with mouse Nanog we used Red/ET based recombineering technique (GeneBridges). Briefly, creERT-pgk-gb2-Hygromycin cassette was inserted into a BAC covering murine Nanog gene region in a way that starting codon of Nanog now became starting codon of CreERT2. Next, BAC fragment containing CreERT insert and appropriate homology arms were cloned into pDTA plasmid encoding Diphtheria Toxin gene cassette used for negative selection in mESCs. BAC recombineered Cre-ER PGK-Hygro construct was linearized with Sca1 and electroporated into the triple allele knock-in mESCs. Hygromycin resistant clones (10 days) were subcloned and screened by southern blot using a 3’ external probe and a PCR creating a 2kb amplification of the 5’ arm, with upstream forward primer and reverse primer from the Nanog Cre-ER knock-in. Correctly targeted clones were transfected with Flippase encoding construct, to remove Hygromycin resistant cassette. Properly targeted mESCs were microinjected into blastocysts to create chimeric animals. At day E13.5 embryos were extracted and used to derive MEFs containing the reporter system with all four knock-in alleles.

### Southern blot analysis

Genomic DNA was extracted from each Hygromycin resistant targeted subclones. 10-15 g of genomic DNA was digested with Mfe1 restriction enzyme for 5 hours and separated by gel electrophoresis. The DNA was transferred to a nitrocellulose membrane that was next hybridized with a radioactive labeled probe and developed using ECL (Thermo).

### iPSC reprogramming by OSKM

For mouse iPSC reprogramming experiments secondary OSKM transgenic reporter MEFs were plated at day 0 in MEF medium: DMEM supplemented with 1% L-Glutamine, 1% Non-essential amino acids, 1% penicillin/streptomycin and 10% FBS, and the following day reprogramming was initiated by changing medium to N2B27 2i/LIF: DMEM-F12 (*BI 01-170-1a*) and Neurobasal medium (*gibco 21103-049*) mixed by 1:1 ratio and supplemented with N2 (*Insulin sigma I-1882, Apo-Transferrin sigma T-1147, Progesterone sigma P8783, Putrescine sigma P5780, sodium selenite sigma S5261*), B27 (*gibco 17504-044*) and 2i: CHIR (*Axon 99021, 3* *M*) PD0325901 (*Tocris 4192, 1* *M*), LIF and adding Doxycycline hyclate (*sigma D9891, 4* *g/mL*), with 4-Hydroxytamoxifen (*sigma H7904, 1 M)* for activation of the Cre-ER reporter system and 5% FBS. Media was replaced every 48 hours, and colonies were analyzed at the indicated time points. Nanog^+/+^ and Nanog^-/-^ ESCs^23^ were rendered transgenic for a constitutively expressed Puromycin resistance cassette^15^, and were subsequently microinjected into host blastocysts. Chimeric embryos were used to derive MEFs, which were selected by Puromycin (2microg/ml final concentration) resistance for at least 3 days then used for transduction and reprogramming in 2i/LIF/15% KSR enriched conditions. In Figure 1a and **Supplementary Figure 1**, MEFs were obtained from Rosa26-m2rtTA (+/-);mCol-OSKM (+/-) mice (Jackson Laboratories #011004), Rosa26-m2rtTA (+/-); mCol STEMCCA-OKSM (+/-) mice (Jackson Laboratories #011001) that were bred with Oct4-GFP or Nanog-GFP knock-in reporter mice 34. NGFP4 iPSC line was generated as previously described^34^.

### Trans-differentiation Reprogramming

Reprogramming of MEFs into cardiomyocytes was done in media containing: DMEM with supplemented 15% FBS and 5% KSR with added JAK1 inhibitor (*Calbiochem 420099*, *0.5* *M*) for 6 days, followed by switch to 1% FBS and 14% KSR and JAK1 inhibitor for 3 days. The medium was supplemented with 1% L-Glutamine, 1% Non-essential amino acids, 1% penicillin/streptomycin and 0.1 mM β-mercaptoethanol. Doxycycline hyclate and 4-Hydroxytamoxifen were added to the medium at day 1 and continuously kept in the medium until day 6-9. From day 9 onwards, cells were cultured in chemically defined RPMI (gibco 21875-034)/N2B27 medium (0.5XN2, 1XB27) with the cardio-inductive growth factor BMP4 (*Peprotech 120-05ET*) for the first 5 days, and supplemented with BSA fraction V 0.05% (*gibco 15260-037*) 1% L-Glutamine, 1% Non-essential amino acids, 1% penicillin/streptomycin and 0.1 mM β-mercaptoethanol. Colonies were allowed to develop for 21 days or until beating cardiomyocytes were noticed among the plated cells.

Reprogramming of MEFS to NSCs was carried out in knock-out DMEM medium (*gibco 10829-018*) with 10% KSR (Invitrogen), 5% FBS, 1% L-Glutamine, 1% Non-essential amino acids, 1% penicillin/streptomycin, 0.05 mM β-mercaptoethanol and with Dox+4OHT for the first 3-6 days. Thereafter, Dox and 4OHT were withdrawn from the medium, and cells were grown in neural reprogramming medium: DMEM-F12 and Neurobasal medium mixed by 1:1 and supplemented with BSA fraction V 0.05%, 1XN2, 1XB27, 1% L-Glutamine, 1% Non-essential amino acids, 1% penicillin/streptomycin and 0.1 mM β-mercaptoethanol with FGF2 (*Peprotech 450-33*, *20ng/mL*) and EGF (*Peprotech 315-09,20ng/mL*), until NSC colonies were formed by day 13. From day 4 onwards JAK1 inhibitor was added to the culture medium. At day 13, formed colonies were subcloned and grown in NSCs medium on Poly-D-Lysine and Laminin coated plates. We note that OSKM activation by Dox for 3 days was sufficient for NSCs trans-differentiation, and 6 days activation for cardiomyocytes trans-differentiation, though substantially fewer colonies developed at these minimally required activation periods as reported in the original papers^10,11^. When applying OSKM-TD protocols without 4OHT or with ETOH as a control replacement, all cardiomyocyte and NSC colonies were negative for tdTomato signal, excluding leakiness of this system under OSKM-TD conditions as similarly observed with iPSCs (Fig. 1D).

Brain-derived NSCs were derived from E13.5 BDF2 mouse embryos according to standard protocols, and were cultured in DMEM/F12 (Invitrogen) supplemented with 1XB27 and 1XN2 cell-supplements (Life Technologies), 0.01% Fraction V BSA 7.5% solution (Life Technologies), 1% Penicillin-streptomycin and 10ng/ml bFGF (R&D) and 20ng/ml EGF (Life Technologies).

### Genetic tracing for X chromosome reactivation in reporter murine female cells

Female *Rosa26 M2RtTa^+/+^* and *m.Col1a-TetO-OSKM^+/+^* mice^18^ *were bred with male m*ice carrying a CMV-GFP knock-in to the X chromosome. Adult Female F1 mice were used as a source for adult tail tip fibroblast or Pro-B somatic cells. GFP-population were sorted on a FACSAria III five-laser equipped cell sorter (BD Biosciences) and further expanded for OSKM-iPSC and OSKM-Td reprogramming experiments as detailed above. Colonies were scored for GFP signal on Zeiss Axioscope D1 microscope, and as expected, iPSCs were homogenously GFP+ (because they maintain both X chromosomes active), while OSKM-Td colonies had patchy GFP expression consistent with random inactivation of the X alleles. Yet the presence of GFP+ cells in TD colonies, indicates reactivation of the X chromosome in the clonal population, followed by random inactivation during differentiation.

### Mouse embryo micromanipulation and imaging

Pluripotent mouse ESCs/iPSCs were injected into BDF2 diploid blastocysts, harvested from hormone primed BDF1 6 week old females. Microinjection into BDF2 E3.5 blastocysts placed in M16 medium under mineral oil was done by a flat-tip microinjection pipette. A controlled number of 10-12 cells were injected into the blastocyst cavity. After injection, blastocysts were returned to KSOM media (Zenith) and placed at 37°C until transferred to recipient females. Ten to fifteen injected blastocysts were transferred to each uterine horn of 2.5 days post coitum pseudo-pregnant females. The latter experiments were approved by Weizmann Institute IACUC (00330111-2).

### Viral production

For primary cell reprogramming, ∼6×10^6^ T-293 cells in a 10 c”m culture dish were transfected with a solution made of 770μL DMEM (Invitrogen) together with 50μl of TransIT^®^-LT1, Δ8.9 (8.5 μg), VSV-G (5.5 μg) and 11μg of the lentiviral target plasmid (FUW-M2rtTA, FUW-TetO-STEMCCA-OKSM, FUW-TetO-OSKM). Viral supernatant was harvested 48 and 72 hours post transfection, filtered through 0.45micron sterile filters (Nalgene) and added freshly to the reprogrammed cells. MSCV-C/EBPα retrovirus stocks were prepared by transient transfection of Phoenix-Eco cells using Fugene (Roche), and supernatants were harvested 48 hours later and filtered. Pro-B cell expansion on OP-9 cells and transduction with C/EBPα was conducted as previously described^34^.

### Immunostaining

Nanog-CreER iPSCs in LIF-free N2B27 medium with supplemented JAK1-Inhibitor and Dox, and *Nanog^-/-^* iPSCs were cultured on glass cover slips (13 mm 1.5H; Marienfeld, 0117530), washed three times with PBS and fixed with 4% paraformaldehyde for 10 minutes at room temperature. Cells were then permeabilized and blocked in 0.1% Triton, 0.1% Tween, and 5% FBS in PBS for 15 min at room temperature. Primary antibodies were incubated for 2 hours at room temperature and then washed with 0.1% Tween and 1% FBS in PBS three times. Next, cells were incubated with secondary antibody for 1 hour at room temperature, washed and counterstained with DAPI, mounted with Shandon Immu-Mount (Thermo Scientific) and imaged.

For staining of trans-differentiated NPCs and cardiomyocytes, cells were reprogrammed in 6 well tissue culture plates, fixed and stained in the tissue culture wells. The following antibodies were used: polyclonal goat Gata4 antibody (*Santa Cruz SC-1237*, *1:200*), mouse monoclonal Myosin antibody (*Hybridoma MF-20*, *1:200*), goat polyclonal Sox1 antibody (*R&D AF-3369*, *1:200*), mouse monoclonal Tuj1 antibody (*Covance MMS-435P*, *1:500*), mouse monoclonal Nestin antibody (*Hybridoma Rat-401*, *1:20*), rabbit polyclonal Sox2 (*Millipore AB5603*, *1:200*), mouse monoclonal Oct4 antibody clone C10 (*Santa Cruz SC5279*, *1:200*), rabbit polyclonal Nanog antibody (*Bethyl A300-397A*, *1:200*) and mouse monoclonal IgM SSEA1antibody (*Hybridoma MC-480 clone*, *1:20*). All antibodies in this study have been validated in the literature and by our internal tests on primary cell lines (data not shown).

### Microscopy imaging

Images were acquired with Axio Scope.A1 microscope (Carl Zeiss) equipped with DP73 camera (Olympus) or with Zeiss Axioscope microscope (Carl Zeiss) equipped with 405 nm, 488 nm, 555 nm and 635 nm solid state lasers, using a ×20 Plan-Apochromat objective (numerical aperture 0.4). All images were acquired in sequential mode. Images were processed with Zeiss Zenblue 2011 software (Carl Zeiss) and Adobe Photoshop CS4.

### Western blot analysis

Whole-cell protein extracts were isolated from wild type Nanog^+/+^ and Nanog^-/-^ ESCs. Blots were incubated with the following antibodies in 5% BSA/TBST: rabbit polyclonal HSP90β (*Calbiochem CA1016*, *1:5,000*), rabbit polyclonal Oct4 antibody clone H-134 (*Santa Cruz SC9081*, *1:1,000*) and rabbit polyclonal Nanog antibody (*Bethyl A300-397A*, *1:1,000*). Secondary antibodies were horseradish peroxidase-linked goat anti-rabbit (1:10,000; Jackson). Blots were developed using ECL (Thermo).

### Teratoma assay

Nanog-CreER iPSCs in LIF-free N2B27 medium with supplemented JAK1-Inhibitor and doxycycline, and *Nanog^-/-^* iPSCs grown in N2B27 2i/LIF cell lines were expended in feeder-free conditions for over 8 passages and injected subcutaneously to the flanks of immune deficient NSG mice. After 4-6 weeks, the mice were sacrificed and the tumor mass extracted and fixed in 4% paraformaldehyde over-night. Slides were prepared from the paraffin embedded fixed tissue, which were next Hematoxylin & Eosin stained and inspected for representation of all three germ layers.

### Alkaline Phosphatase staining

Cells were grown on 6 well tissue culture plates and were washed three times with PBS and fixed with 4% paraformaldehyde for 10 minutes at room temperature. Alkaline phosphatase staining was performed according to the manufacturer protocol (*Millipore SCR004*).

### RNA sequencing and gene expression profiling analysis

RNA was extracted from Trizol pellets, and utilized for RNA-seq by TruSeq RNA Sample Preparation Kit v2 (Illumina) according to manufacturer’s instruction. DNA sequencing was conducted on Illumina Hiseq1500. For sample alignment and processing, Tophat software version 2.0.10 was used to align reads to mouse mm10 reference genome (UCSC, December 2011). Read counts per exon were generated using coverageBed function, from BEDTools version 2.16.2, for all exons in mm10 ensemble GTF (UCSC, December 2011). In this process the read counts were calculated for 628,052 exons. Exons were normalized to RPKM values, by dividing read counts per exon by the exon length in Kbp and number of aligned reads per sample in million reads. To further filter, only exons that have at least one call higher than 2 were included resulting in 371,807 exons corresponding to 19,755 genes. Gene expression was defined by the expression (RPKM) level of the highest expressed exon associated to this gene. For gene expression analysis, Matlab version R2011b was used. Sample clustering with all 19,755 genes **(Supplementary Fig. 3c)** was done with hierarchical clustering using Spearman correlation as a distance metric and average linkage. The hierarchical clustering utilize a normalization procedure over gene expression, where each gene is normalized to have zero mean and standard deviation equal to 1 over the samples, i.e., each row is normalized.

### Primer Sequences

M2RtTa genotyping:

Forward primer 5’ AAA GTC GCT CTG AGT TGT TAT 3’

Reverse primer 5’ GCG AAG AGT TTG TCC TCA ACC 3’

OSKM Col1a genotyping:

Forward primer 5’ GCA CAG CAT TGC GGA CAT G 3’

Reverse primer 5’ TTG CTC AGC GGT GCT GTC CA 3’

tdTomato:

Forward primer 5’ CTG TTC CTG TAC GGC ATG G 3’

Reverse primer 5’ GGC ATT AAA GCA GCG TAT CC 3’

CreER:

Forward primer 5’TGG ATG TGG AAT GTG TGC CAG 3’

Reverse primer 5’CCC TAG TGG CTT CCA AAT TCA 3’

